# Synchronized infection identifies early rate-limiting steps in the hepatitis B virus life cycle

**DOI:** 10.1101/2020.04.05.026237

**Authors:** Anindita Chakraborty, Chunkyu Ko, Christin Henning, Aaron Lucko, Xiaodong Zhuang, Jochen M. Wettengel, Ulrike Protzer, Jane A McKeating

**Affiliations:** Institute of Virology, Technical University of Munich, School of Medicine /Helmholtz Zentrum München, Munich, Germany; Technical University of Munich, Institute for Advanced Study, Munich, Germany; Nuffield Department of Medicine, University of Oxford, Oxford, UK; German Center for Infection Research (DZIF), Munich, Germany

## Abstract

Hepatitis B virus (HBV) is an enveloped DNA virus that contains a partially double-stranded relaxed circular (rc) DNA. Upon infection, rcDNA is delivered to the nucleus where it is repaired to covalently closed circular (ccc) DNA that serves as the transcription template for all viral RNAs. Our understanding of HBV particle entry dynamics and host pathways regulating intracellular virus trafficking and cccDNA formation is limited. The discovery of sodium taurocholate co-transporting peptide (NTCP) as the primary receptor allows studies on these early steps in viral life cycle. We employed a synchronized infection protocol to quantify HBV entry kinetics. HBV attachment to cells at 4°C is independent of NTCP, however, subsequent particle uptake is NTCP-dependent and reaches saturation at 12h post-infection. HBV uptake is clathrin- and dynamin dependent with actin and tubulin playing a role in the first 6h of infection. Cellular fractionation studies demonstrate HBV DNA in the nucleus within 6h of infection and cccDNA was first detected at 24h post-infection. Our studies show the majority (83%) of cell bound particles enter HepG2-NTCP cells, however, only a minority (<1%) of intracellular rcDNA was converted to cccDNA, highlighting this as a rate-limiting in establishing infection *in vitro*. This knowledge highlights the deficiencies in our *in vitro* cell culture systems and will inform the design and evaluation of physiologically relevant models that support efficient HBV replication.

## INTRODUCTION

Hepatitis B Virus (HBV) infects 257 million individuals worldwide and is a major driver of end-stage liver disease, cirrhosis, and hepatocellular carcinoma (HCC). HBV is an enveloped DNA and prototypic member of the *hepadnaviridae* that establishes its genome as an episomal, covalently closed circular DNA (cccDNA) in the nucleus of infected hepatocytes. Current treatments suppress viral replication but are not curative, largely due to the persistence of the cccDNA transcriptional template and failure to mount an effective anti-viral immune response^1^. In most cases, treatment is life-long and patients may still develop HCC^2^, highlighting a clinical need for new curative therapies^3^. Despite its central role in the HBV life cycle our understanding of the host factors regulating cccDNA genesis and half-life is limited^4^.

Viral entry into a host cell represents the first step in the infectious life cycle and is mediated via specific interactions between virus encoded proteins and cellular receptors that define internalization pathways^5^. The discovery that sodium taurocholate co-transporting polypeptide (NTCP) acts as a receptor for HBV^6,7^ enabled the development of *in vitro* culture systems that support the complete HBV replication cycle. HBV encodes three envelope glycoproteins, small (S), middle (M) and large(L)^8^. The preS1 domain of the L protein binds heparan sulfate proteoglycan (HSPG)^9–11^ that precedes high-affinity virus interaction with NTCP. Synthetic peptides mimicking the preS1 binding site for NTCP, such as Myrcludex-B (MyrB) inhibit HBV infection^12,13^ and a recent phase II clinical trial efficacy in HBV patients co-infected with hepatitis delta virus^14^. To date the role NTCP plays in HBV internalization is not well defined and the virus has been reported to use both clathrin and caveolin-dependent endocytic pathways^15–18^. HBV engagement of NTCP was recently shown to activate Epidermal Growth Factor receptor and this signalling pathway promotes virus translocation to the endosomes via undefined pathways^19^. Our understanding of the host pathways regulating HBV uptake and intracellular particle trafficking is limited and warrants further investigation.

Current HBV culture systems use high viral inocula (ranging from 500 to 10,000 genome equivalents of HBV per cell) and frequently require polyethylene glycol (PEG)-mediated precipitation to initiate infection^20–23^, suggesting that our *in vitro* model systems are inefficient and may not recapitulate the liver environment. Asabe *et al*^24^ reported that a single HBV particle was sufficient to infect a chimpanzee, illustrating the infectious nature of HBV particles in vivo. To explore the early steps in the HBV life cycle required to infect human hepatocyte derived cells expressing NTCP we established a synchronized infection protocol to quantify HBV internalization and early intracellular trafficking events. Our studies show a relatively efficient process of HBV internalization and particle trafficking to the nucleus with >80% of cell-surface attached virus entering permissive cells. However, the conversion of newly imported partially double-stranded relaxed circular DNA (rcDNA) to cccDNA was inefficient, uncovering a rate-limiting step in establishing productive infection of current *in vitro* model systems.

## RESULTS

### Quantifying HBV attachment and internalization

To quantify HBV attachment and internalization kinetics we used a well-established method^25,26^ where virus is allowed to bind to cells on ice, cultures shifted to 37°C to promote viral uptake and non-internalized virus removed with trypsin (**Fig.1**). This protocol enables a synchronized uptake of HBV particles into target cells that can be enumerated by PCR quantification of HBV rcDNA genomes. HBV stocks for these experiments were purified on a heparin affinity column, followed by sucrose gradient centrifugation, and include mature type-B infectious HBV particles^27^ and L-containing filamentous sub-viral particles. Polyethylene glycol 8000 (PEG) is routinely used to enhance HBV infection in cell culture systems including primary human hepatocytes, HepaRG cells and more recently HepG2-NTCP cells^20–23^. Since this agent precipitates virus and has been reported to promote herpes simplex virus type 1 fusion at the plasma membrane^28^ and Semliki forest virus (SFV) infection of non-permissive cell types^29^ all experiments were conducted without PEG. We previously reported that HepG2 cells engineered to express NTCP^21^ support HBV replication and we selected the K7 sub-clone for our kinetic experiments and confirmed NTCP expression (**Fig.2a**). Our initial experiments optimized the trypsinization protocol to ensure removal of cell-associated non-internalized virus (**Fig.2b**). We noted a comparable dose-dependent binding of HBV to HepG2 and HepG2-NTCP cells (**Fig.2b-c**), demonstrating that viral attachment at this low temperature is independent of NTCP.

**Fig.1:**
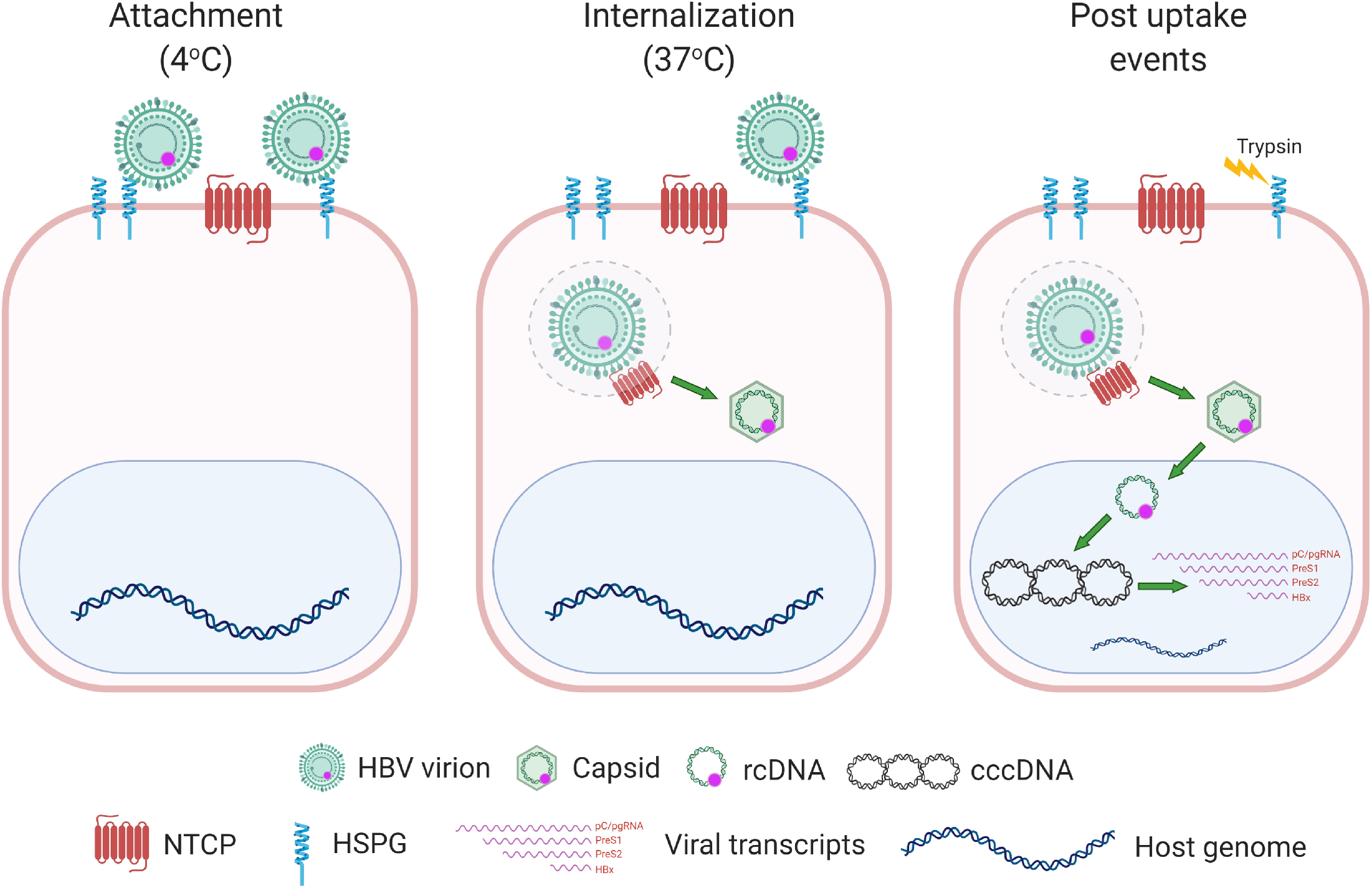
Cartoon depicting a synchronized HBV infection protocol. Pre-chilled cells were inoculated with HBV for 1h and cells moved to 37°C, leading to synchronous internalization of viral particles. At various times the cells are treated with trypsin to remove cell-associated non-internalized viral particles and viral parameters quantified.

**Fig.2:**
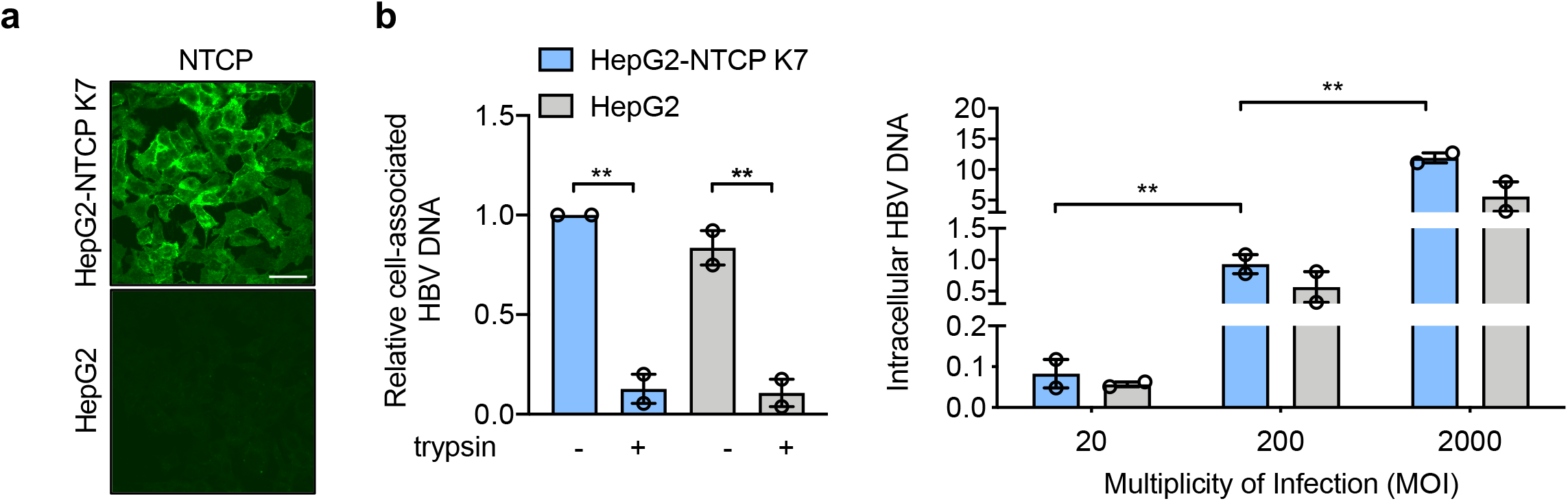
Quantifying HBV attachment to target cells. (**A**) *HBV attachment to HepG2 cells is NTCP independent*. HepG2 and HepG2-NTCP K7 were stained for NTCP expression using Atto488 labelled Myrcludex B (200 nM) and imaged using 63x objective (scale bars indicate 20 μm). Pre-chilled HepG2 and HepG2-NTCP K7 cells were inoculated with HBV (MOI 200) for 1h on ice, unbound virus removed by washing and cells treated with trypsin or left untreated and cell-associated HBV DNA quantified by RT-PCR. Data is expressed relative untreated HepG2-NTCP cells. (**B**) *HBV attachment to HepG2 cells is dose-dependent*. Increasing dose of HBV (MOI 20 – 2000) was inoculated with HepG2-NTCP K7 and HepG2 cells for 1h at 4°C, unbound virus removed by washing and cell-associated HBV DNA quantified. HBV DNA levels are expressed relative to *PRNP* and represent two independent experiments presented as mean ± SEM. Each experiment consisted of three replicates per condition. Statistical analysis was performed using a Mann–Whitney *U* test (*p<0.05, **p<0.01, ***p<0.001).

To assess the role of NTCP in HBV internalization we inoculated HepG2 and HepG2-NTCP cells with virus (multiplicity of infection, MOI, of 200 genome equivalents per cell) for 1h at 4°C and transferred the cultures to 37°C for 1h, 3h or 6h prior to digesting with trypsin to remove non-internalized particles and quantifying cell-associated HBV DNA. We observed a time-dependent increase in trypsin-resistant HBV DNA after culturing the cells at 37°C and noted significantly higher levels of viral DNA in HepG2-NTCP after 6h compared to HepG2 cells (**Fig.3a**), showing a clear role for NTCP in HBV internalization. We confirmed these observations with human Huh-7 hepatoma cells that showed similar kinetics and NTCP-dependency of HBV internalization (**Fig.3a**). The majority of *in vitro* studies on HBV replication utilize dimethyl sulfoxide (DMSO) to arrest target cell proliferation and we were interested to evaluate the effect of DMSO on HBV internalization. We noted comparable levels of internalized HBV DNA in DMSO-treated and untreated HepG2-NTCP cells (**Supp Fig.1**), demonstrating a minimal role for DMSO in modulating early steps in viral uptake.

**Fig.3:**
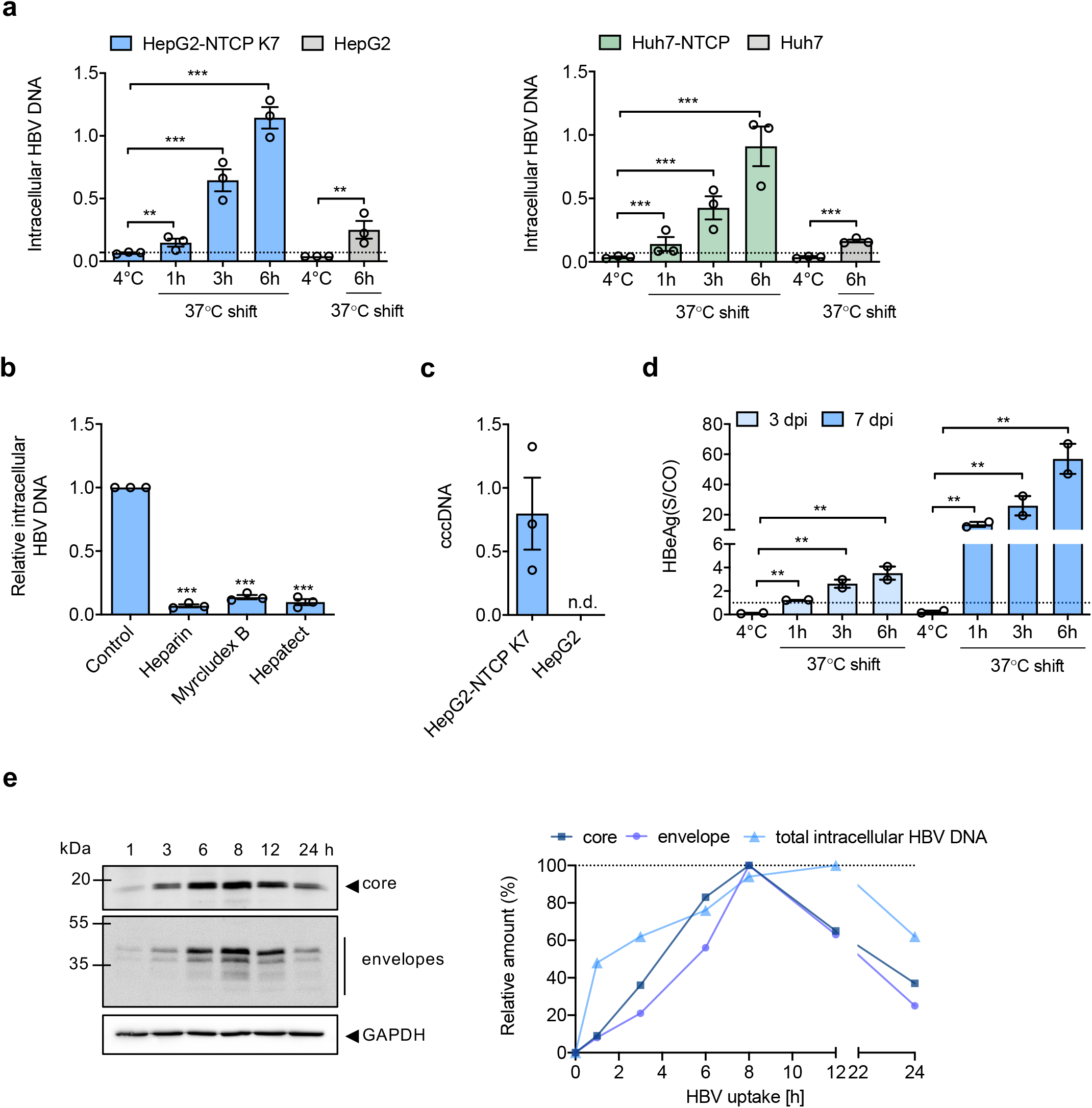
HBV internalization kinetics. (**A**) *HBV internalization is temperature and NTCP-dependent*. HepG2 and Huh-7 hepatoma cells and those engineered to express NTCP were inoculated with HBV (MOI 200) and trypsinized after 1h at 4°C or following incubation at 37°C for 1, 3 or 6h. Intracellular HBV DNA levels are expressed relative to *PRNP* and the dotted line represents trypsinized 4°C samples that were set as background for the assay. (**B**) *Receptor and HBV glycoprotein dependent particle internalization*. HepG2-NTCP K7 cells were inoculated with HBV (MOI 200) in the presence or absence of heparin (50 IU/ml), Myrcludex B (200 nM) or Hepatect (0.5 IU/ml) and trypsin-resistant intracellular HBV DNA copies measured after 6h. Data is expressed relative untreated HepG2-NTCP cells. (**C**) *Short-term synchronized HBV infection of HepG2-NTCP cells generates cccDNA*. Parental HepG2 and HepG2-NTCP K7 cells were inoculated with HBV (MOI 200) as detailed above and after 6h at 37°C cells were typsinized and cultured at 37°C for 3d before measuring cccDNA. HBV cccDNA levels are expressed relative to *PRNP* and represent two independent experiments presented as mean ± SEM. (**D**) *Association between internalized HBV particles and HBeAg expression*. HepG2-NTCP K7 were inoculated with HBV (MOI 200) and trypsinized after 1h at 4°C or following incubation at 37°C for 1, 3 or 6h and the infected cells cultured for 3 or 6d before measuring extracellular HBeAg. Dotted line represents the limit of detection of the assay, where all values above 1 are considered positive. (**E**) *HBV internalization kinetics*. HepG2-NTCP K7 cells were inoculated with HBV (MOI 200) as detailed above (A) and after defined times at 37°C the trypsinized cells were lysed and probed for HBV envelope and core proteins by western blot and images quantified by densitometry. A summary of internalization kinetics is depicted as the amount of intracellular HBV DNA, core or envelope proteins and plotted as relative data where the highest value of the respective parameter is set to 100%. Data are representative of up to three independent experiments presented as mean ± SEM. Each experiment consisted of three replicates per condition. Statistical analysis was performed using a Mann–Whitney *U*test (*p<0.05, **p<0.01, ***p<0.001), n.d: not detected.

To assess the specificity of our viral internalization assay we evaluated the effect of known HBV entry inhibitors: heparin that competes for virus attachment to cellular HSPGs^10^; MyrB^30^ and Hepatect a polyclonal anti-HBV Ig that neutralizes viral infection^31^. All of the entry inhibitors were used at a concentration that neutralized >90% of HBV infection (judged by HBeAg expression at 5d post-infection) and significantly reduced intracellular HBV DNA levels by more than 80% after 6h post inoculation (**Fig.3b**). These data confirm that HBV internalization is dependent on cellular HSPGs, NTCP and viral surface glycoproteins.

Since we noted a low level of HBV internalization into HepG2 cells we were interested to know if this virus could establish a productive infection and cultured the cells at 37°C for 3 days and monitored cccDNA levels. We only detected cccDNA in HepG2-NTCP cells, demonstrating non-productive uptake pathway(s) in the parental HepG2 cells that lack NTCP (**Fig.3c**). To determine whether early HBV internalization steps limit productive infection, we cultured HepG2-NTCP cells following 1h, 3h or 6h synchronized infection for 3 and 7 days and measured HBeAg expression as a marker of cccDNA transcriptional activity. It is noteworthy that HBV particles are trypsin sensitive and show an approximate 10-fold reduction in infectivity. We noted a significant association between HBeAg levels and inoculation time that persisted after 3 and 7 days of culture (**Fig.3d**), demonstrating that the amount of internalized virus defines virus replication.

To extend and validate these observations we assessed HBV internalization kinetics in HepG2-NTCP cells by monitoring HBV particle associated core or surface envelope glycoproteins **(Fig.3e)**. Densitometric scanning of western blots showed a peak of intracellular core and surface glycoprotein-associated particles at 8h and a subsequent decline over the duration of the assay. Intracellular HBV DNA showed a delayed peak at 12h post internalization. Given the semi-quantitative nature of western blots, these data are in good agreement with earlier PCR data and show a time-dependent internalization of HBV particles that reaches saturation at 8-12h post inoculation.

### HBV entry kinetics in dHepaRG cells

The bipotent HepaRG cell line can be differentiated toward biliary-like epithelial cells and hepatocyte-like cells that express endogenous NTCP and support HBV replication^32^. In our experience we generally observe a 1:1 ratio of hepatocyte:biliary cells and staining the differentiated cells for NTCP expression using MyrB showed a low frequency of NTCP expressing cells compared to HepG2-NTCP cells (**Supp Fig.2a**). Attempts to study HBV uptake into dHepaRG cells in the absence of PEG yielded negligible results with no detectable intracellular viral DNA. We previously reported that including PEG 8000 in the inoculum increased HBV uptake 10-fold^21^ and inoculating dHepaRG cells with HBV in the presence of PEG showed comparable viral uptake kinetics over a 6h period to HepG2 and Huh-7 cells over-expressing NTCP (**Supp Fig.2b**). These data suggest that NTCP expression levels *per se* have a negligible impact on the kinetics of HBV internalization but may regulate the absolute levels of internalized virus. Our attempts to study HBV internalization into PHHs yielded poor quality data with limited evidence of viral uptake even in the in the presence of PEG and showed high donor variability.

### Cellular pathways regulating HBV internalization

To study the cellular pathways that regulate HBV internalization we evaluated a panel of pharmacological agents that target various cellular trafficking pathways: Dynasore inhibits dynamin and arrests vesicle formation from the plasma membrane^33^; Pitstop 2 inhibits clathrin-mediated endocytosis^34^ and ethyl-isopropyl amiloride (EIPA) targets the Na^+^/H^+^exchanger, inhibiting macropinocytosis^35,36^. We confirmed the specificity of Pitstop and Dynasore in targeting clathrin- and dynamin-dependent endocytosis in HepG2-NTCP cells by their inhibition of FITC-labelled transferrin uptake and Vesicular Stomatitis virus (VSV) pseudoparticle infection (**Supp Fig.3**). All agents were evaluated for their ability to inhibit HBV internalization into HepG2-NTCP or Huh-7 NTCP cells after 6h inoculation and HBeAg expression measured after 5 days. Heparin and MyrB were included as positive controls known to inhibit HBV uptake. Pre-treating cells with Dynasore and Pitstop and maintaining compounds during the viral inoculation stage inhibited HBV uptake into both cell lines (**Fig.4a-b**), suggesting a dynamin and clathrin dependent endocytic uptake process. In contrast, EIPA, had no effect on HBV internalization, suggesting a negligible role for micropinocytosis in viral uptake into hepatoma cells. These observations were validated by measuring HBeAg expression 5 days post inoculation (**Fig.4a-b**). To study the role of the host cytoskeleton in regulating HBV uptake, HepG2-NTCP cells were treated with either nocodazole or cytochalasin D that interfere with microtubule and actin dynamics, respectively^37,38^ **(Fig.4c).** Both of these agents significantly inhibited HBV uptake in the first 6h following infection and HBeAg levels, suggesting a role for microtubule and actin filaments in regulating intracellular capsid trafficking.

**Fig.4:**
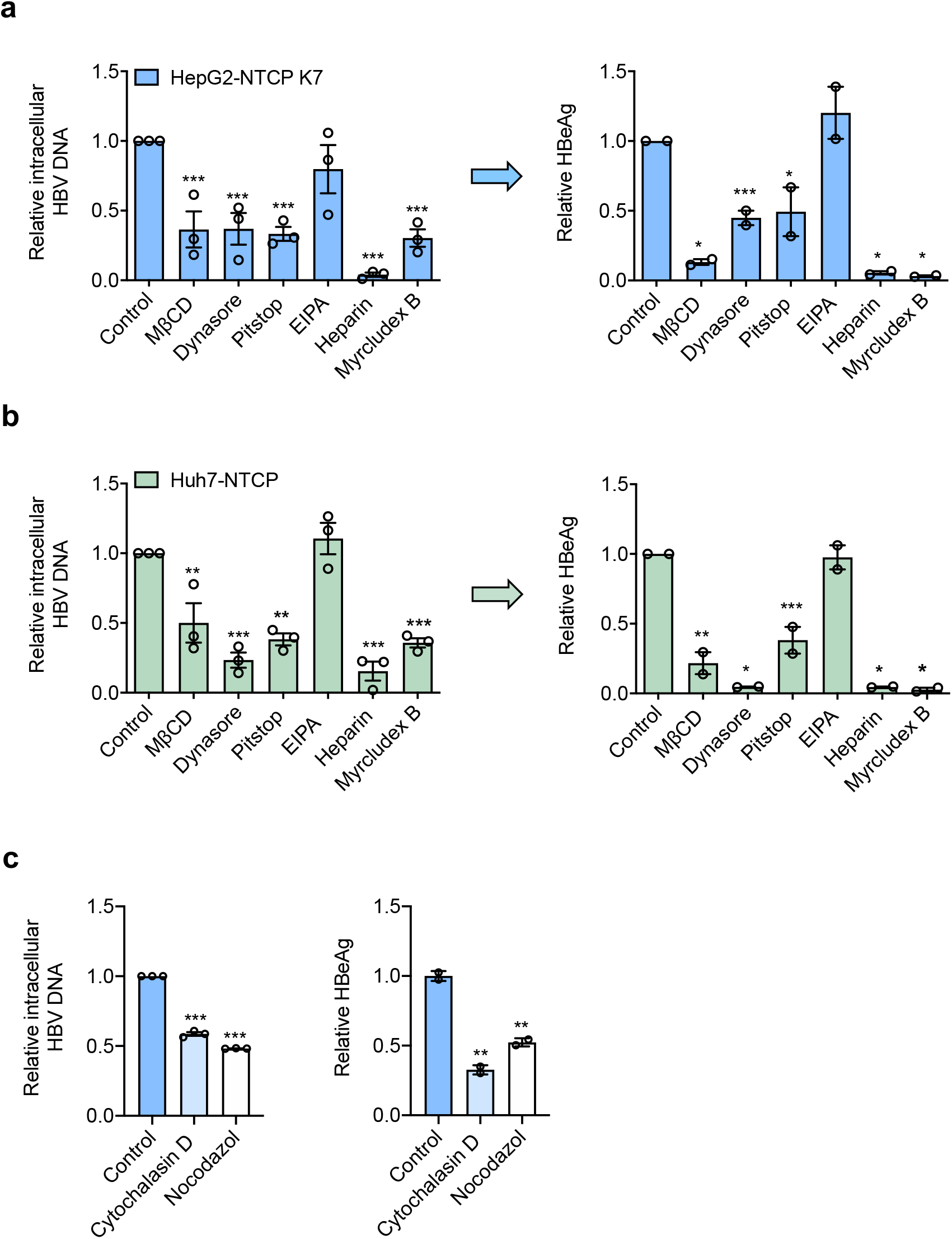
Cellular trafficking pathways exploited by HBV. *HBV internalization is clathrin and dynamin dependent*. HepG2-NTCP K7 (**A**) or Huh7-NTCP (**B**) cells were treated with pharmacological agents targeting dynamin-(Dynasore: 100 μM), clathrin-mediated endocytosis (Pitstop: 50 μM) or micropinocytosis (EIPA: 100 μM) and inoculated with HBV (MOI 200) as detailed in Fig.1. Cells were pre-treated with Dynasore and Pitstop for 0.5h prior to infection and during the HBV inoculation step. EIPA was co-treated during HBV inoculation. Trypsin-resistant intracellular HBV DNA copies after 6h or extracellular HBeAg expression after 5d was measured. Data are plotted relative to untreated control and represent up to three independent experiments presented as mean ± SEM. (**C**) *HBV internalization is actin and tubulin dependent*. HepG2-NTCP cells were treated with actin and tubulin disrupting agents, Cytochalasin D and Nocodazole (50μM each) respectively and inoculated with HBV (MOI 200). Trypsin-resistant intracellular HBV DNA after 6h or extracellular HBeAg levels after 5d were measured. Data are plotted relative to untreated control and represent up to three independent experiments presented as mean ± SEM. Each experiment consisted of three replicates per condition. Statistical analysis was performed using a Mann–Whitney *U* test (*p<0.05, **p<0.01, ***p<0.001).

### Kinetics of HBV trafficking from the cytoplasm to the nucleus

To study the early steps in the HBV life cycle that precede cccDNA genesis we analyzed subcellular fractions for intracellular HBV DNA. A synchronized HBV infection was performed and cytoplasmic and nuclear fractions harvested at early (1, 3, 6, 8 and 12h) and late (24, 48 and 72h) time points post-trypsinization for quantification of HBV DNA and cccDNA. Cellular fractionation was confirmed by probing cytoplasmic and nuclear samples for α-tubulin and lamin A/C, respectively and DNA samples for presence of the housekeeping gene *PRNP* (**Suppl Fig.4**). Intracellular HBV DNA was first detected in the cytoplasm within 1h of incubating the cells at 37°C and particle trafficking to the nucleus was detected after 3h (**Fig.5a)**. HBV DNA levels in the nucleus and cytoplasm were saturated by 12h and we noted 3.5-fold higher levels of viral DNA in the cytoplasm compared to the nucleus as well as a loss of viral DNA in both nuclear and cytoplasmic fractions after 24h (**Fig.5a**). We used published PCR methodologies^21^ to quantify cccDNA in the nuclear and cytoplasmic fractions and first detected cccDNA in the nuclear fraction 24h post-infection, which subsequently increased throughout the duration of the experiment (**Fig.5b**).

**Fig.5:**
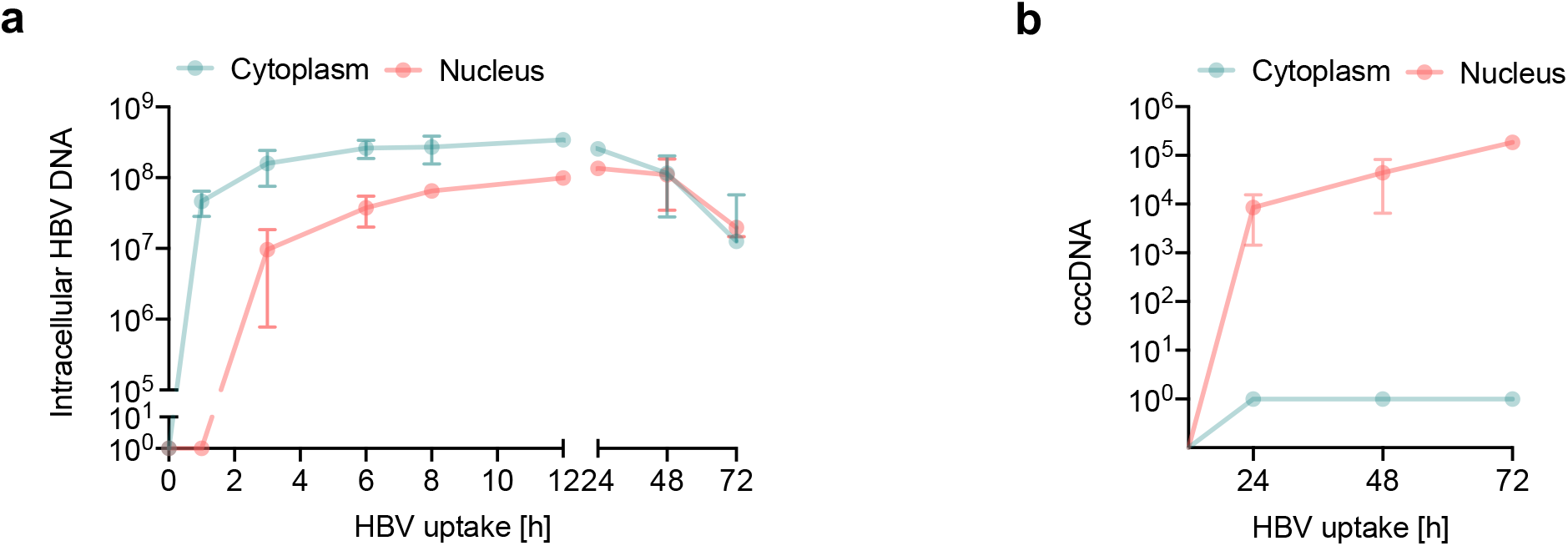
Kinetics of HBV trafficking to the nucleus. (**A**) *Intracellular HBV trafficking in cytoplasm and nucleus*. Synchronised HBV infection (MOI 200) where cytoplasmic and nuclear samples were harvested at the indicated times. HBV DNA was first detected in cytoplasm at 1h and in the nucleus after 3h and monitored thereafter. Detection threshold of the qPCR lies between 10-100 copies. (**B**) *Synchronized infection and cccDNA levels*. HepG2-NTCP K7 cells were inoculated with HBV (MOI 200) as detailed above (A) and cccDNA levels in the cytoplasmic and nuclear fraction measured. Absolute numbers of total intracellular HBV DNA and cccDNA in cells/cm^2^ are shown. Data are representative of two independent experiments presented as mean ± SEM. Each experiment consisted of duplicates per condition. Statistical analysis was performed using a Mann–Whitney *U* test (*p<0.05, **p<0.01, ***p<0.001).

### Identifying rate-limiting steps in HBV infection

Having optimized the synchronized infection protocol, we quantified HBV attachment (1h at 4°C), internalization (6h, the inferred half-maximal value) and cccDNA (72h) levels in HepG2 and HepG2-NTCP cells. Similar levels of virus inocula attached to HepG2-NTCP and HepG2 at 4°C (25% and 17%, respectively), consistent with a role for HSPGs in defining the initial association of virus with the cell surface (**Table 1**). The majority (84%) of cell-bound particles entered HepG2-NTCP cells. We observed a surprisingly high level (46%) of intracellular HBV DNA in HepG2 cells and since we failed to detect any cccDNA in these cells, this most likely reflects a non-productive uptake pathway in these cells (**Table 1**). Finally, we noted that less than 1% of the intracellular HBV DNA detected at 6h was converted into cccDNA by 72h. It is worth noting that the detection limit of our assays for quantifying rc- and cccDNA is 100 copies per reaction, suggesting that our earlier conclusion is not biased by differences in the sensitivity of the PCR methods. In summary, these data show that particle internalization is efficient with the majority of cell-bound particles entering NTCP expressing cells, with at least 22% of particle associated DNA reaching the nucleus within 12h. In contrast, the subsequent conversion of incoming rcDNA to cccDNA is inefficient, identifying a rate-limiting step in establishing productive infection.

**Table 1:** Rate limiting steps in the early HBV life cycle. Summary of synchronized infection in HepG2-NTCP and HepG2 cells. Data showing input HBV DNA of the inoculum, cell-associated HBV DNA (attachment) and ntracellular HBV DNA after 6h post infection (p.i.) at 37°C along with cccDNA levels at 72h p.i. Absolute copy numbers are presented from cells/cm^2^. These data are presented as the mean ± SEM of eight independent experiment with biological duplicates. n.d: not detected.

|  | HepG2-NTCP K7 |  |  | HepG2 |  |  |
| --- | --- | --- | --- | --- | --- | --- |
|  | Absolute copy number<br>per cm<sup>2</sup> | % from<br>input virus | % from<br>attached<br>virus | Absolute copy number<br>per cm<sup>2</sup> | % from<br>input virus | % from<br>attached<br>virus |
| HBV inoculum | $1.5 \times 10^8 \pm 2.9 \times 10^7$ | - | - | $1.5 \times 10^8 \pm 2.9 \times 10^7$ | - | - |
| Attachment | $3.7 \times 10^7 \pm 4.7 \times 10^6$ | $25 \pm 6$ | - | $2.6 \times 10^7 \pm 4.1 \times 10^6$ | $17 \pm 5$ | - |
| Total intracellular<br>HBV DNA (6h p.i.) | $3.1 \times 10^7 \pm 2.5 \times 10^6$ | $21 \pm 6$ | $84 \pm 21$ | $1.2 \times 10^7 \pm 1.9 \times 10^6$ | $8 \pm 3$ | $46 \pm 17$ |
| cccDNA (72h p.i.) | $2.2 \times 10^5 \pm 3.8 \times 10^4$ | $0.15 \pm 0.05$ | $0.6 \pm 0.16$ | - | - | - |

## DISCUSSION

Our current knowledge on the early steps of HBV infection are not well defined, despite their key role in determining tissue and species tropism. To address this gap in our knowledge we developed an assay to quantify particle internalization and nuclear transport to identify rate-limiting steps in the early viral life cycle. HBV showed comparable binding to HepG2 and HepG2-NTCP chilled on ice suggesting that primary attachment events are independent of NTCP, consistent with a role for HSPG in mediating low affinity attachment of HBV to target cells^10,39^. A recent report using recombinant HBV particles confirmed the HSPG-dependency of particle attachment and suggested an intracellular role for NTCP^40^.

Our kinetic studies show a clear role for NTCP in regulating HBV uptake into HepG2 and Huh-7 cells and for establishing a productive infection. We noted a time-dependent increase in particle uptake quantified by measuring intracellular HBV DNA or virus associated core and envelope proteins that saturated after 12h. The time for viral DNA and proteins to reach their maximum levels may reflect differential half-lives of the genomic material and protein. HBV DNA was first detected in the cytoplasm after 1h and in the nuclear fraction by 3h. HBV core protein encodes a nuclear localization sequence that targets capsids to the nuclear pore complex (NPC) in an importin a/ ß mediated manner^41–43^, however, these studies did not address the time for HBV capsids to reach the NPC. Similar entry kinetics were reported for duck hepatitis B infection (DHBV) of primary duck hepatocytes (PDH), showing DHBV DNA in the nucleus by 4h^44^. Importantly, these data are consistent with reports of intracellular trafficking times for other enveloped viruses including HIV that can target the nucleus within 4h of infection^45,46^.

Recent advances in imaging technologies can visualise the internalization of virus particles (reviewed in^47^). Several studies have reported the use of fluorescent labelled HBV viral structural proteins or sub-viral particles^17,48^.

However, these approaches come with certain constraints as the labelling can impair viral infectivity and do not always resemble natural infection. Our attempts to image HBV large (L) envelope protein of internalized virus provided very weak signals in first 3h of infection. Combinatorial approaches to visualise HBV envelope and genome may provide a better understanding of viral fusion, uncoating and nuclear transport. Herrscher *et al* recently reported HBV in clathrin-coated pits and vesicles using electron microscopy (EM) and cryo-EM with immunogold labelling^18^ consistent with a clathrin-dependent endocytic entry route.

We first detect cccDNA in the synchronized infection assay after 24h, consistent with reports for DHBV infection^44^. Since our PCR method to quantify cccDNA uses a T5 exonuclease to remove non-cccDNA species, this treatment may result in a loss of >20% of cccDNA^21^. Given these caveats our data suggests that a minority of intracellular encapsidated rcDNA (<1%) is converted to cccDNA. The slow conversion of rcDNA to cccDNA may simply reflect the rate of genome uncoating and trafficking across the nuclear membrane. However, our fractionation studies showing that up to 22% of total intracellular DNA is in the nucleus within the first 6-12h of infection suggeststhat nuclear targeting is not rate limiting for cccDNA genesis. The mechanism of rcDNA conversion to cccDNA is not fully defined and a number of host pathways have been reported^49–52^. The viral polymerase is removed by tyrosyl DNA phosphodiesterase 2 (Tdp2)^7,53,54^ and this is followed by the removal of the RNA primer by a cellular flap-like structure specific endonuclease, Fen1^50,55^. Host Polymerase k and DNA ligase (Lig) I and III^56^ complete the cccDNA synthesis^57^. All of these factors may be limiting in human hepatoma cells and may contribute to the inefficient rcDNA to cccDNA conversion.

Our results support a role for a clathrin and dynamin in defining HBV particle uptake into HepG2-NTCP and Huh-7-NTCP cells, consistent with a recent report assessing these pathways in HBV infection^18^. In contrast, EIPA had no detectable effect on HBV uptake, suggesting a minimal role for caveolin-dependent micropinocytosis. This contrasts to observations reported by Macovei et al^15^ showing a role for caveolin-1 in HBV infection of dHepaRG cells. These differences may be the result of infection protocols, where PEG-enhanced infection may promote viral aggregation and non-physiological uptake pathways. In addition, we and others^18^ noted low to undetectable levels of caveolin-1 expression in HepG2-NTCP cells and PHHs compared to dHepaRG cells, which may also contribute to the different entry pathways reported in these studies. In contrast to many other enveloped viruses that enter cells via clathrin mediated endocytosis, HBV uptake kinetics is slower, for example VSV and SFV require only several minutes to enter their target cell and establish infection^58,59^.

In summary, our studies show the majority of cell bound particles enter NTCP expressing target cells, however, only a minority of intracellular rcDNA is converted to cccDNA, highlighting this as a rate-limiting step in establishing infection *in vitro*. We believe this knowledge is essential to aid the interpretation of mechanistic studies probing pathways regulating cccDNA genesis and half-life and for screening anti-viral agents. Furthermore, this data will inform the design of physiologically relevant models that support efficient HBV replication.

## EXPERIMENTAL METHODS

### Cell lines

HepG2-NTCP K7 cells and Huh-7 NTCP cells^21,60^ were cultured in Dulbecco’s Modified Eagles Medium F12 (DMEM-F12) supplemented with 10% fetal bovine serum (FBS) and penicillin/streptomycin. HepaRG cells were cultured in Williams E medium supplemented with 10% FBS, 50 U penicillin/streptomycin mL^-1^, 5 μg human insulin mL^-1^ and 5×10^-7^ M hydrocortisone hemisuccinate (Sigma). Cells were seeded and expanded for two weeks and subsequently differentiated for another two weeks in the presence of 1.8% DMSO^61^ (dHepaRG).

### HBV purification protocol

HBV was purified and concentrated from HepAD38 cell culture supernatant using previously published protocols^27,62^. In brief, cells were cultivated in multi-layer flasks and supernatant collected every 3-4 days. Collected supernatant was loaded on 5 mL Heparin HiTrap columns and eluted with 390 mM NaCl. Subsequently eluent was loaded on top of a layered sucrose gradient (3mL 60%, 7mL 25% and 9mL 15%) in a SW32Ti centrifugation tube. After 3.5h centrifugation at 32.000 rpm in a SW32Ti rotor, the gradient was fractionated in 2 mL aliquots and the virus-rich fraction (2nd from the bottom) titrated, aliquoted and stored at −80°C.

### Synchronized HBV infection

Cells were seeded on collagen-coated plates and differentiated for two days with media containing 2.5% DMSO. Cells were pre-chilled on ice for 15 minutes and cold HBV containing inoculum added to cells on ice for 1h enabling the virus to bind to the cell surface. Medium was exchanged and cells were subsequently shifted to 37 °C for 1-72h. For harvesting cells were washed with PBS and trypsinized for 3 minutes. Samples were either subjected to DNA extraction for HBV DNA or cccDNA analysis or samples collected for HBeAg measurement.

### Quantification of HBV DNA and cccDNA by real time PCR

Total cellular DNA was extracted using a NucleoSpin Tissue kit (Macherey Nagel) according to the manufacturer’s protocol. Intracellular HBV DNA and cccDNA was analysed as previously described in^21^. For cccDNA analysis total DNA was subjected to T5 exonuclease digestion. Total intracellular HBV DNA (HBV1844For: 5’-GTTGCCCGTTTGTCCTCTAATTC-3’ and HBV1745Rev: 5’- GGAGGGATACATAG-AGGTTCCTTGA-3’) and cccDNA (cccDNA92For: 5’-GCCTATTGATTGGAAAGTATGT-3’ and cccDNA2251Rev: 5’-AGCTGAGGCGGTATCTA-3’) were normalised to human prion protein *PRNP* (PRNPFor:5’- TGCTGGGAAGTGCCATGAG-3’ and PRNPRev:5’-CGGTGCATGTTTTCACGATAGTA-3’). An external plasmid standard was used for absolute quantifications.

### HBeAg ELISA

HBeAg was qualitatively quantified using an automated BEP III system (Siemens Healthcare).

### NTCP staining of cell surface

Cells were seeded on collagen-coated plates. 200 nM Atto488-labeled Myrcludex B was added for 30 min at 37°C on cells.^7,63^ Images were obtained by fluorescence microscopy.

### Western blotting

Samples were lysed with Pierce RIPA buffer (Thermo Scientific Fisher) supplemented with a protease inhibitor cocktail ^64^ on ice for 10 min. 4x Laemlli buffer was added to obtain a final 1x concentration and incubated at 95°C for 5 min. Proteins were separated on a 12% polyacrylamide gel and transferred to PVDF membranes (Amersham). The membranes were blocked in PBST, 3% milk (Sigma). Primary antibody detecting, core (in house supernatant 8C9) and envelope protein (H863, provided by H.Schaller), lamin A/C (Thermo Scientific Fisher), GAPDH (Acris) and a-tubulin (Sigma) were incubated in 1% milk overnight. Proteins were detected using a HRP coupled secondary antibody with the Amersham ECL Prime Western Blotting Detection Reagent.

### Subcellular fractionation

Samples were separated using NE-PER™ Nuclear and Cytoplasmic Extraction Reagents (Thermo Scientific Fischer) according to manufacturer’s protocol. Intracellular DNA from cytoplasm or nucleus was extracted using NucleoSpin Tissue kit (Macherey Nagel) and qPCR for HBV DNA and cccDNA performed.

## ACKNOWLEDGEMENTS

We thank Peter Wing for critical reading of the manuscript and Stephan Urban (University of Heidelberg) for his generous provision of Myrcludex B.

## AUTHOR CONTRIBUTIONS

AC designed and conducted experiments and co-wrote the manuscript; CK designed experiments and co-wrote the manuscript, CH conducted experiments; XZ conducted an experiment; JW provided reagents; UP designed the study and co-wrote the manuscript, JAM designed the study and co-wrote the manuscript.

