## Supplemental figures 1 - 4 for "Synchronized infection identifies early rate-limiting steps in the hepatitis B virus life cycle"

Supplementary Figures

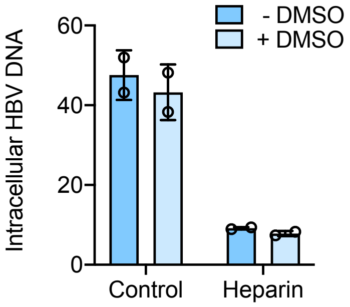


**Supplementary Fig. 1: Effect of DMSO on HBV entry.** HepG2-NTCP K7 cells either pre-treated with 2.5% DMSO for 3d (+DMSO) or untreated (-DMSO) were inoculated with HBV (MOI, 200) for 1h on ice and then incubated at 37°C for 6h, trypsinized and intracellular HBV DNA quantified and expressed relative to PRNP. Heparin (50 IU/ml) was included as control to inhibit HBV uptake.

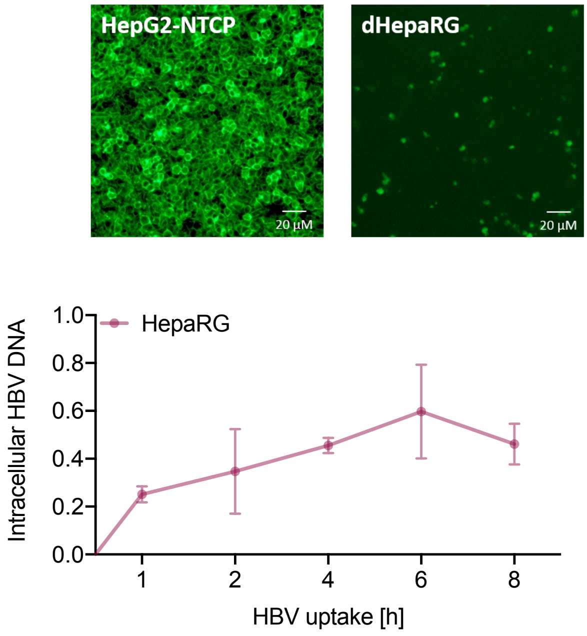


**Supplementary Fig.2: HBV internalization into dHepaRG cells. (A)** HepG2-NTCP cells and dHepaRG cells were stained with uAtto488 labelled Myrcludex B (200 nM)) and fluorescent images acquired using a Zeiss fluorescence microscopy. (**B**) HepaRG cells were inoculated with HBV (MOI 200) in the presence of 4% PEG 800 for 1h on ice, unbound virus removed by washing and analysed at indicated time points (0-8h) for total intracellular HBV DNA. Data is expressed relative to *PRNP*.

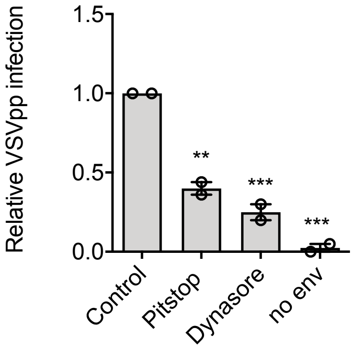


**Supplementary Fig.3: Effect of pharmacological agents on VSV-G infection**. HepG2-NTCP K7 cells were pre-treated with Dynasore and Pitstop for 0.5h prior to infection and during VSVpp inoculation for 24h. Pseudoparticles lacking an envelope protein were included as a negative control. Data are representative of two independent experiments presented as mean ± SEM. Each experiment consisted of duplicates per condition. Statistical analysis was performed using a Mann–Whitney *U* test (*p<0.05, **p<0.01, ***p<0.001).

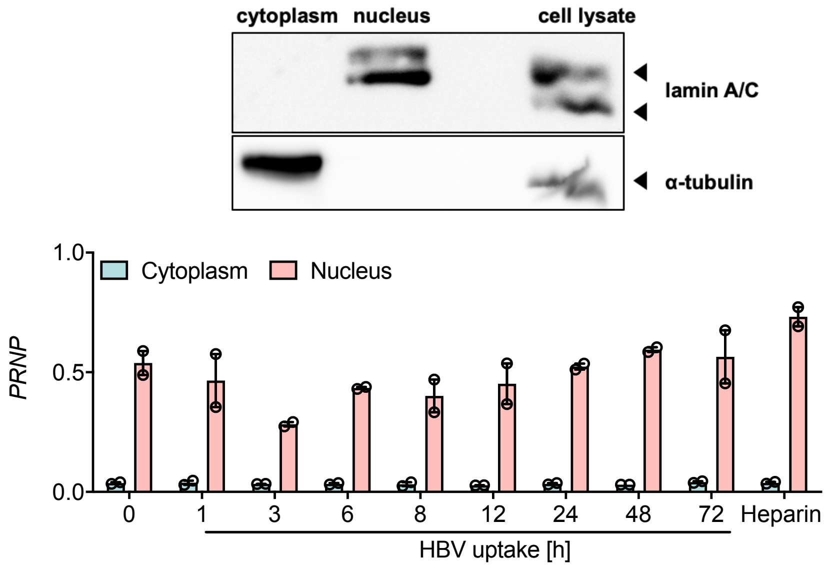


**Supplementary Fig.4: Subcellular fractionation. (A)** Western blot targeting nuclear and cytoplasmic resident protein, lamin A/C and α-tubulin, respectively. Whole cell lysate was included as a control. **(B)** Synchronised HBV entry was assessed and samples harvested at indicated time points (0-72h) were subjected to qPCR analysis of cytoplasmic and nuclear extracted DNA for *PRNP*. Data are representative of two independent experiments presented as mean ± SEM. Each experiment consisted of duplicates per condition.
